# Training Data Diversity Enhances the Basecalling of Novel RNA Modification-Induced Nanopore Sequencing Readouts

**DOI:** 10.1101/2024.08.29.610342

**Authors:** Ziyuan Wang, Ziyang Liu, Yinshan Fang, Hao Helen Zhang, Xiaoxiao Sun, Ning Hao, Jianwen Que, Hongxu Ding

## Abstract

Accurately basecalling sequence backbones in the presence of nucleotide modifications remains a substantial challenge in nanopore sequencing bioinformatics. It has been extensively demonstrated that state-of-the-art basecallers are less compatible with modification-induced sequencing signals. A precise basecalling, on the other hand, serves as the prerequisite for virtually all the downstream analyses. Here, we report that basecallers exposed to diverse training modifications gain the generalizability to analyze novel modifications. With synthesized oligos as the model system, we precisely basecall various out-of-sample RNA modifications. From the representation learning perspective, we attribute this generalizability to basecaller representation space expanded by diverse training modifications. Taken together, we conclude increasing the training data diversity as a novel paradigm for building modification-tolerant nanopore sequencing basecallers.

## INTRODUCTION

During nanopore sequencing, biomolecules with various chemical structures translocate through protein pores further produce squiggling electric signals^1^. While such a rationale promises the routine detection of nucleotide chemical modifications, these modifications bring substantial challenges for accurately basecalling underlying sequence backbones. State-of-the-art basecallers, such as Guppy, Bonito and Dorado delivered by the Oxford Nanopore Technologies, were built upon deep neural networks^2^. Extensive studies have reported that such basecallers are susceptible to modifications, which commonly exist in native DNA/RNA molecules. Most importantly, modification-induced basecalling artifacts are systematic, which cannot be resolved by simply increasing the sequencing depth as what we in general do for random errors. Indeed, these non-random errors can serve as informative signatures for determining well-studied modifications^3-8^. The large number of uncharacterized modifications further aggravate this problem: it is widely-believed that a majority of biologically-relevant nucleotide modifications remain undiscovered^9,10^; among the >50 DNA^11^ and >170 RNA^12^ modifications that have been discovered *in vivo*, most of them have yet to be nanopore sequenced. Unlike the well-studied modifications, without prior biological knowledge, we are agnostic about the types and locations of basecalling errors produced by uncharacterized modifications. Taken together, these systematic and in most cases “cryptic” artifacts significantly undermine the basecalling accuracy.

The precise basecalling, on the other hand, serves as the prerequisite for virtually all the downstream bioinformatic analyses, including genome assembly and structural variation characterization, transcriptomic alternative splicing and expression quantification as well as DNA/RNA modification detection^13^. To better accomplish such biological applications, basecallers that are agnostic of modifications, especially the previously-uncharacterized ones, are therefore in pressing need. Here, we present a novel paradigm for basecalling previously-unseen modifications, by combining diverse existing modifications as training data. Specifically, the latest deep learning basecallers in general consist of encoder and decoder neural networks. The encoder network will condense sequencing signals into a highly-informative representation space. Diverse training modifications can expand such a space, to a degree that out-of-sample modifications can also be properly encoded. By this means, previously-unseen modifications will be precisely basecalled by the decoder network, as shown in Figure 1.

**Figure 1.**
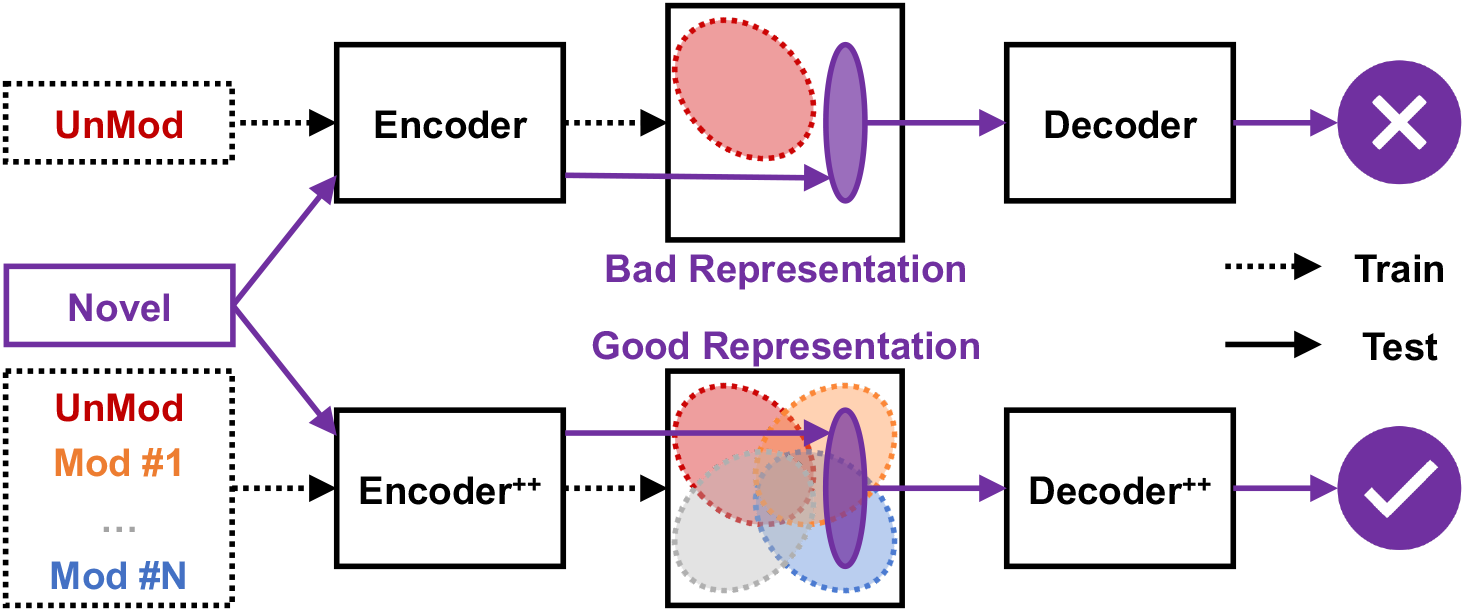
Overview of the basecaller training strategy. Basecallers in general consist of encoder and decoder neural networks: encoders first condense nanopore sequencing readouts as highly-informative representations; decoders further transform the produced representations as nucleotide sequences. Diverse training modifications will expand the representation space, thus making basecallers generalizable to novel modifications. UnMod and Mod denote unmodified and modified training data categories, respectively. Novel denotes the out-of-sample modification in the test data. Encoder^++^ and Decoder^++^ comprise the basecaller trained with diverse modifications, as opposed to the basecaller trained with only the UnMod data.

## RESULTS

### An oligo-based model system for investigating nanopore sequencing basecalling

Generic basecallers, which in theory could handle any biological and artificial nucleotide sequences, are extremely compute-intensive and data-demanding to train. We therefore leverage control oligos as the model system to develop and evaluate basecallers. In line with previous studies^3,6,14^, our model system contains 4 oligo backbones, which together covered all 1,024 RNA 5mers with a median occurrence of 10. These diverse sequence contexts were adopted in order to ensure the soundness of our basecalling analyses. To explore the basecalling of modification-induced nanopore sequencing readouts, besides unmodified (UM) oligos, 8 additional modified derivatives including N1-methyladenosine (m1A), N6-methyladenosine (m6A), N4-acetylcytidine (ac4C), 5-methylcytosine (m5C), 5-hydroxymethylcytosine (hm5C), 5-methyluridine (m5U), pseudouridine (Psi) as well as N1-methylpseudouridine (m1Psi), were collected for our analyses (see METHODS).

### Unmodified sequences are inadequate to train the modification basecaller

We first examined whether a basecaller trained using only unmodified nucleotide sequences can properly handle their modified counterparts. We therefore trained a basecaller using UM oligos, then executed it on UM, m1A, m6A, ac4C, m5C, hm5C, m5U, Psi and m1Psi test oligos. We quantified the basecalling accuracy “functionally” with downstream alignment CIGAR (see METHODS). As the positive control, UM oligos were accurately basecalled with a 99.80% average match rate, which confirmed the high-quality basecaller training. We also noticed that m5C, hm5C and m5U oligos were acceptably basecalled (99.48%, 98.57% and 99.18% average match rate, respectively), which suggested the UM-trained basecaller could be generalized to a limited number of modifications. The remaining test groups, in particular ac4C, Psi and m1Psi, drastically decreased basecalling confidence and produced considerably more basecalling errors (Figure 2A). For instance, we found an average 2.23%, 6.45% and 8.42% increase in deletion, which was the most common basecalling error in our analysis, for ac4C, Psi and m1Psi compared to UM respectively. Taken together, such results demonstrated that basecallers trained with only unmodified sequences are less generalizable to analyze modification-induced nanopore readouts.

**Figure 2.**
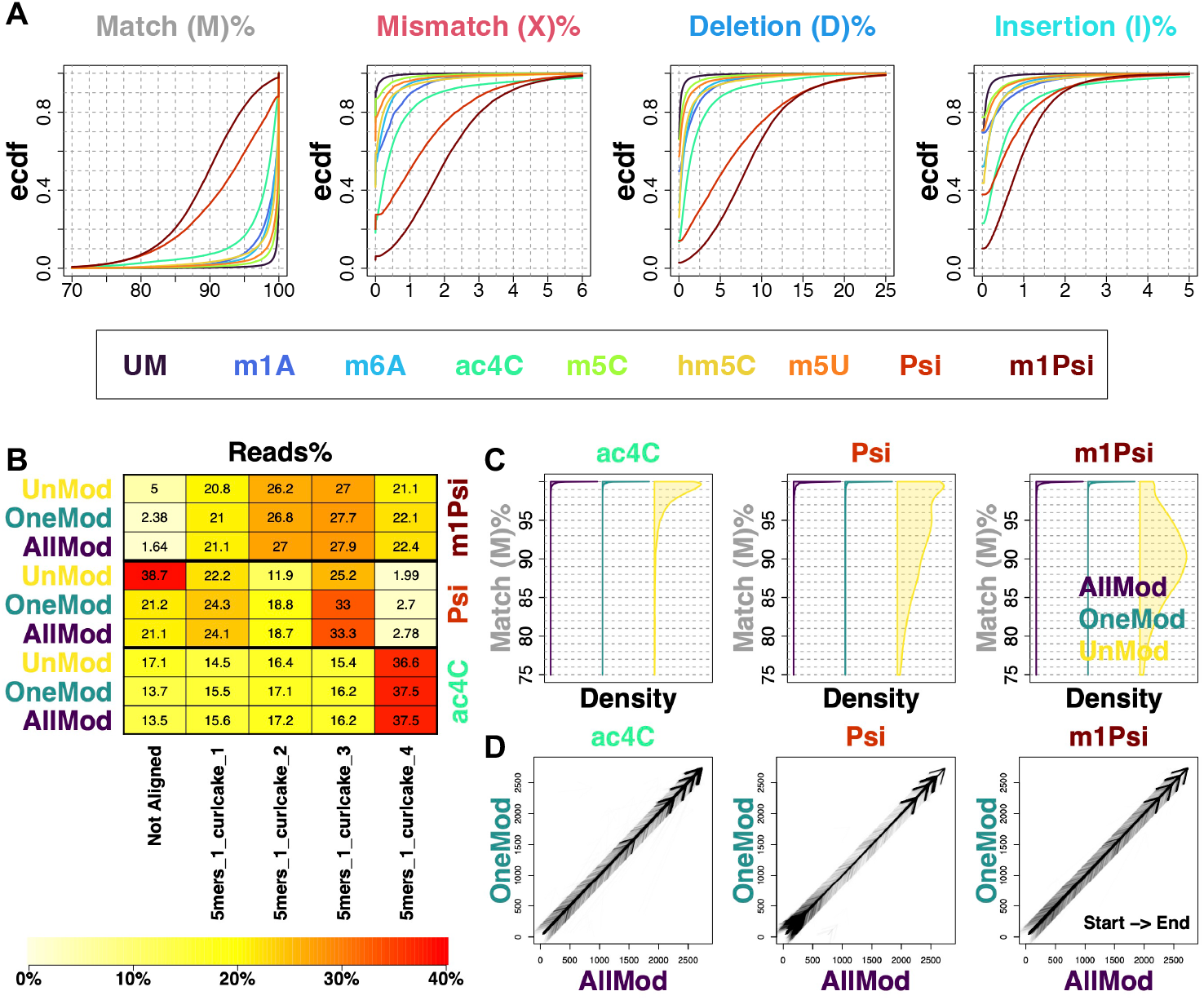
Basecallers trained with diverse known modifications gain the capability to basecall novel modifications. (A) Performance of the basecaller trained only by the unmodified data on all the read groups. Basecalling performance was assessed with the per-read CIGAR alignment fraction, including match (M), mismatch (X), deletion (D) and insertion (I). UM and acronyms stand for unmodified and modified RNA oligo categories, respectively. Ecdf denotes the empirical cumulative distribution function. Performance of basecallers trained by combining all the oligo groups except for ac4C, Psi or m1Psi was quantified. Specifically, the mappability (B) and per-read CIGAR match fraction (C) were used as quantification metrics. AllMod, the basecaller trained by all the modifications except for the one to be basecalled; OneMod, the basecaller trained with only the modification to be basecalled; UnMod, the basecaller trained by only unmodified reads. (D) Alignment position comparison between “AllMod” and “OneMod”. Start and End denote the alignment direction.

### Diverse training modifications improve the basecalling of novel modifications

We next asked whether combining diverse known modifications in the training dataset could facilitate the basecalling of novel modifications. We therefore investigated ac4C, Psi and m1Psi, which were identified as “least analyzable” modifications with the UM-basecaller. Specifically, we trained corresponding basecallers with all other modifications except for the ones to be analyzed (the “AllMod” groups). Meanwhile, we trained basecallers using only candidate modifications (the “OneMod” groups) as positive controls. We also used the above UM-basecaller (the “UnMod” groups) to evaluate the baseline basecalling performance (see METHODS). As shown in Figure 2B, “AllMod” generated comparable mappability with “OneMod” positive controls for all the three modification types, with absolute differences <0.8%. As shown in Figure 2C, “AllMod” increased basecalling accuracy compared to “UnMod”: we noticed an average increase of 2.60%, 7.30% and 10.40% in CIGAR matches for ac4C, Psi and m1Psi, respectively. Most importantly, “AllMod” basecalling accuracy reached the same level as “OneMod” positive controls: we noticed a negligible <0.40% absolute difference for all the three oligo types. Finally, as shown in Figure 2D, the “AllMod” training will polish, rather than biasing basecalling, by generating consistent alignment positions compared to the “OneMod” ground-truth.

In summary, our results demonstrated that increasing training modification diversity will enhance the basecalling performance of out-of-sample novel modifications. We further confirmed such a conclusion with the latest Bonito and Dorado basecalling systems, as shown in Figure S1.

### Evaluating the out-of-sample basecalling generalizability for training modification combinations

We further asked, are all the training modifications required for accurate out-of-sample basecalling? To formally answer such a question, an evaluation metric for training modification combinations is required. Inspired by our discovery that basecallers trained with only unmodified oligos basecalled certain modifications (m5C, hm5C, m5U) with acceptable accuracy (Figure 2A), as well as signals of such modifications were less deviated from their canonical counterpart (Figure S2, see METHODS), we hypothesized that the optimal training combinations can completely cover signals of the out-of-sample modification. We therefore defined the “signal cover score”, with the following five steps: 1) correspond signal segments (events) with basecalled sequences; 2) assemble all the events mapped to the same sequence position, calculate their signal mean values, then use 10% and 90% quantiles (to avoid outliers) as the “effective signal range”; 3) quantify the training effective signal range, by taking the union of all the training modifications; 4) measure the cover fraction, as the test effective signal range that can be covered by the training counterpart; 5) calculate the final signal cover score, by taking the summation of all the cover fractions along the candidate sequence (Figure 3A).

**Figure 3.**
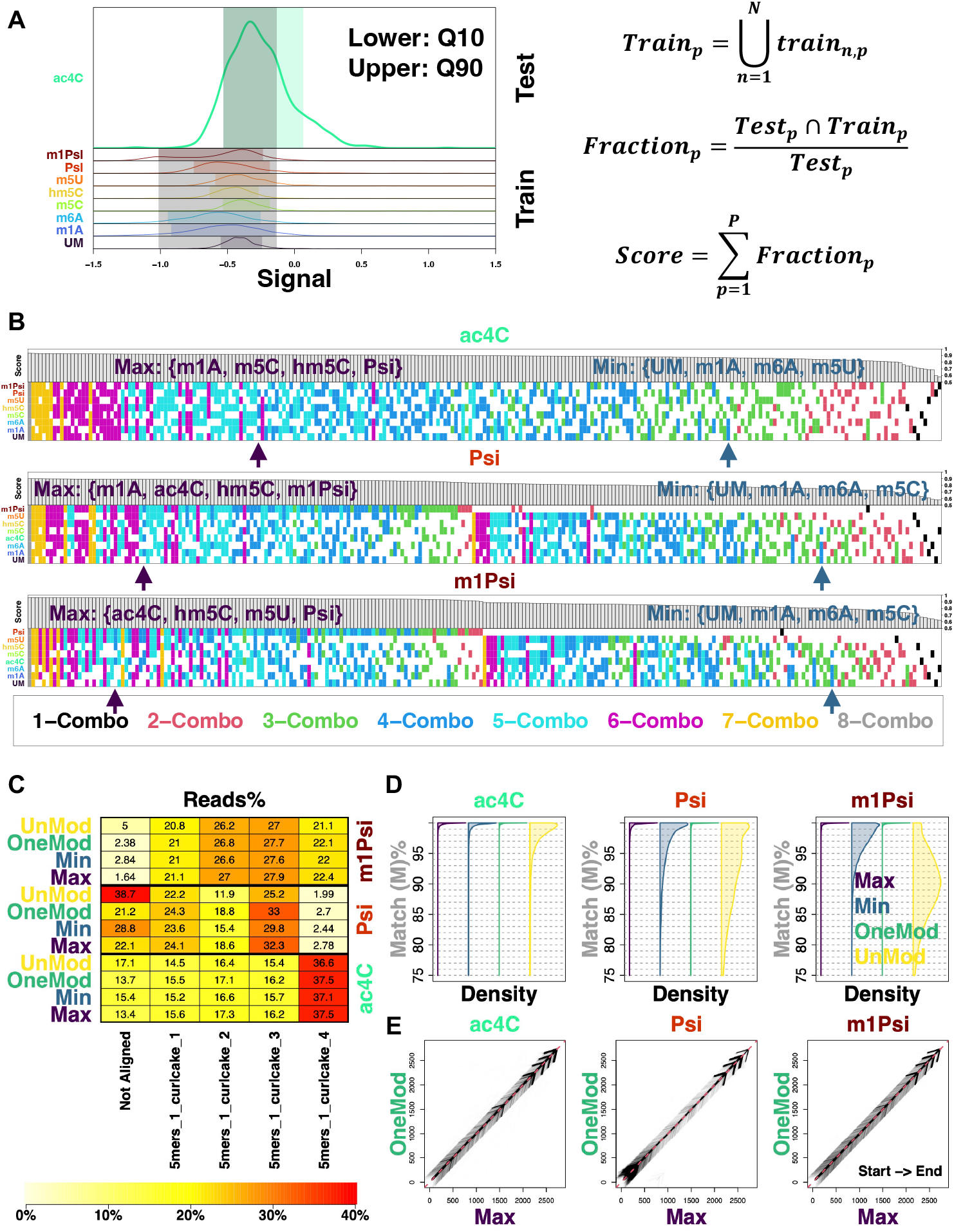
Prioritizing training combinations for precise out-of-sample modification basecalling using signal cover scores. (A) The definition of “signal cover score”. Q10 and Q90 mark the 10% and 90% signal quantiles, respectively. *N* and *P* denote the total number of training modification classes and sequence positions, respectively. (B) Signal cover scores for all the possible training modification combinations in descending order, for ac4C, Psi and m1Psi test groups. Bars denote the inclusion of training modifications, and corresponding colors represent numbers of modifications, for a certain combination. Specifically, 4-combo combinations with the highest (Max) and lowest (Min) signal cover scores were marked. The performance of Max and Min basecallers was next quantified. Specifically, the mappability (C) and per-read CIGAR match fraction (D) were used as quantification metrics. AllMod, the basecaller trained by all the modifications except for the one to be basecalled; UnMod, the basecaller trained by only unmodified reads. (E) Alignment position comparison between “AllMod” and “Max”. Start and End denote the alignment direction.

Based on such a signal cover score, for ac4C, Psi and m1Psi test groups, we evaluated all the possible training modification combinations. As shown in Figure 3B, we observed that combinations of more modifications in general produced higher signal cover scores, which echoes with our observation that diverse training data improves the basecalling of out-of-sample modifications. In particular, we noticed that the inclusion of Psi and m1Psi in training modifications significantly improved signal cover scores of m1Psi and Psi test data, respectively. These findings echo with our results that Psi and m1Psi reads can be acceptably handled by basecallers training using only m1Psi and Psi reads, respectively (Figure S4A and S5A). We further systematically confirmed that signal cover scores can be adopted to evaluate training modification combinations. Without losing generality, we prioritized the analyses of 4-combo combinations, and highlighted ones with the highest (Max) and lowest (Min) scores. We found that “Max” generated comparable mappability and basecalling accuracy with “OneMod” positive controls (basecallers trained with only candidate modifications), and out-performed “Min”, particularly in Psi and m1Psi groups, as shown in Figure 3C and D. We finally confirmed that “Max” could generate consistent alignment with “OneMod” ground-truth, as shown in Figure 3E. Therefore, we concluded the signal cover score as the metric for prioritizing training modification combinations.

### Precise basecalling requires high-quality data representations

We then related the basecaller accuracy with the quality of its encoder representation space. Representation learning condenses neural network inputs into a highly-informative representation space to achieve downstream tasks^15^. During basecalling, nanopore sequencing signals will be encoded in the representation space then decoded as nucleotide sequences. To explore how data representations affect the basecalling accuracy, we analyzed ac4C test oligos. In particular, we trained the “All” basecaller by combining all the oligo categories (except for ac4C), as well as 8 single-category (without ac4C) basecallers (see METHODS).

We observed that compared to the individually-trained basecallers, “All” can significantly promote the CIGAR match fraction. Such precise basecalling was consistently observed in different regions among all the four oligos. We further observed that although artifacts made by individually-trained basecallers were in general prevalent, certain regions were more likely to be accurately analyzed. For example, “m1A” decently analyzed the region 800 to 820 of the first oligo, which was highlighted with the red box (Figure 4A). We next investigated such region-specific elevated basecalling performance, by interrogating the encoder representation space. Without losing generality, we prioritized the above boxed region, by projecting corresponding nanopore signals into the representation space (see METHODS). Within the representation space, we quantified the read-level similarity and found that, in the more accurate “UM”, “m1A”, “m6A” and “hm5C” basecallers, ac4C test reads were more similar to their corresponding training reads. On the contrary, test-train similarity was significantly reduced among error-prone “m5C”, “m5U”, “Psi” and “m1Psi” basecallers. Most importantly, the highest test-train similarity was produced by the most precise “All” basecaller (Figure S3A). We further visualized such a representation space similarity pattern between test and training reads with UMAP, as shown in Figure 4B. In representation learning, test data points can be properly encoded, if and only if they fall in the manifold produced by training data points. We therefore highlight the importance of a high-quality representation space in precise basecalling. We further confirmed this conclusion with Psi (Figure S3B, S4A-C) and m1Psi (Figure S3C, S5A-C) test oligos.

**Figure 4.**
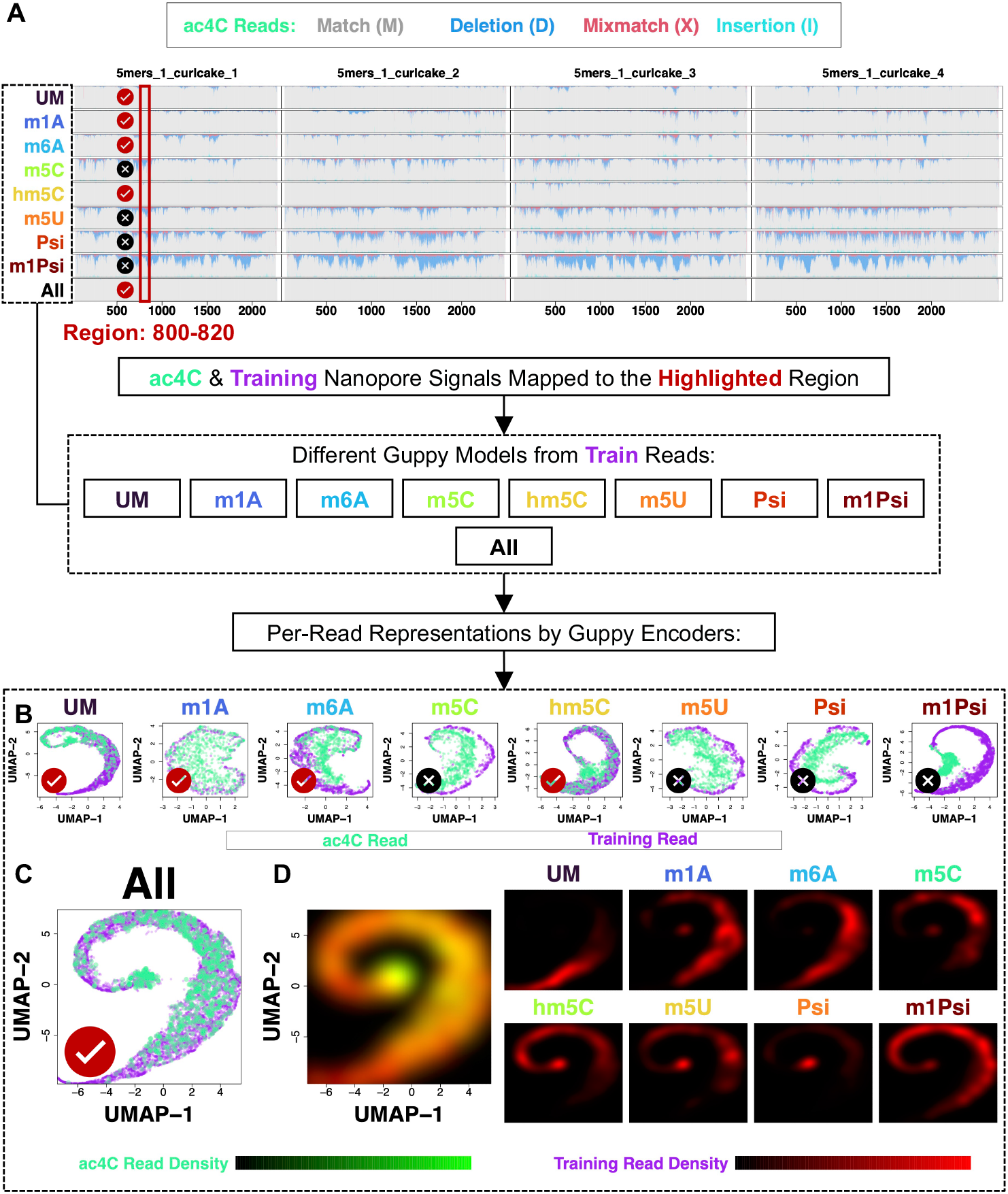
Diverse training data expands the representation space thus making the basecaller generalizable to novel modifications. (A) Performance of individually and jointly-trained basecallers on ac4C reads was visualized with the genome viewer graph, which shows per-nucleotide CIGAR fractions. All, the jointly-trained basecaller by all the oligo types except for ac4C; other acronyms denote individually-trained basecallers. For individually (B) and jointly-trained (C) basecallers, read fragments mapped to the boxed region were first converted as representation vectors with different basecaller encoders, then visualized by a UMAP plot. Train denotes reads used for training the corresponding basecaller. (D) Spatial distributions of different oligo types in the UMAP space as shown in (C). Black-to-green and red palette denotes ac4C and training reads, respectively.

### Training data diversity yields a generalizable basecaller representation space

We further explained, from a representation learning perspective, that diverse training oligos will expand the representation space, to a degree that out-of-sample novel modifications could also be properly encoded. Specifically, we revisited the “All” representation space, and assessed the spatial distribution of ac4C, as well as different types of training oligos (Figure 4D). We presented the spatial density of ac4C and training data points using the black-to-green and black-to-red palette, respectively. We first confirmed that the majority of ac4C points were covered with the training manifold, by finding negligible stand-alone green area. We further found that the entire training manifold was required to thoroughly encode ac4C data, by finding most representation space to be yellowish. We also found that training groups occupied different sub-space, which together completed the training manifold. Taken together, these results suggested that diverse training modifications will complement each other, thereby generalizing the representation space to out-of-sample modifications. We further confirmed such a conclusion with Psi (Figure S4D) and m1Psi (Figure S5D) test oligos.

## DISCUSSION

Promoting the basecalling accuracy, especially for native DNA/RNA sequencing signals, remains a central challenge in nanopore sequencing bioinformatics. This is because the latest basecallers are less tolerant to modifications, which commonly exist among native DNA and RNA molecules and will deviate nanopore sequencing signals (Figure S2). To properly address this limitation, basecallers that are agnostic to nucleotide modifications are therefore in urgent need.

Here, we presented a novel paradigm for training such modification-tolerant basecallers: we demonstrated that previously-unseen modifications could be precisely basecalled by including diverse existing modifications in training data. We anticipated such a paradigm increasing the basecalling accuracy of native DNA/RNA nanopore sequencing readouts, further shedding light on diverse biological usecases, e.g. *de novo* genome assembly by generating highly-accurate DNA contigs^16^, mRNA vaccine quality analyses by rigorously assessing sequence, length, integrity and purity^17^, and DNA/RNA modification detection by delivering precise backbones to scrutinize modification status of each nucleotides^18^.

To fully leverage such a paradigm, we emphasized the quality and generalizability of the basecaller representation space. In particular, a high-quality representation space could extract all essential information for a precise basecalling; a generalizable representation space could tolerate sequencing signal variations for the novel modification basecalling. To optimize the basecaller representation space, as the potential future direction, we will leverage self-supervised learning techniques^19^ for training foundation model encoders^20^. During the self-supervised learning process, the encoder neural network first transforms inputs into a representation space. The decoder neural network then reconstructs inputs from the encoded representation space. The recreation of inputs suggests the negligible information loss in the representation space. By this means, all the information harbored inside nanopore sequencing signals could be extracted by the encoder, further providing a solid foundation for the precise basecalling. Foundation models are trained by a broad range and excessive amount of data, therefore can be applied across diverse usecases. This paradigm echoes with our discovery, that diverse training oligos will expand, further generalize the representation space for basecalling various out-of-sample modifications. Altogether, we expect the foundation model encoder trained via self-supervised learning to be able to produce the high-quality and generalizable representation space, therefore enhancing the performance of modification-tolerant basecalling.

## METHODS

### RNA Oligo Synthesis

RNA oligo sequences (“curlcakes”) were reported in ^3^, and were cloned into engineered pUC57 *in vitro* transcription (IVT) plasmids. We then incorporated modified nucleotides into curlcakes with IVT. We produced unmodified and m6A RNA oligos for resequencing because existing datasets^3^ were sequenced over 5 years ago and may be outdated. We also included m1A, which was not surveyed by previous studies^3,6,15^, in our modification collection. Specifically, curlcake-containing pUC57 plasmids were digested using EcoRV and BamHI restriction enzymes for at least two hours at 37 °C, and further analyzed with agarose gel electrophoresis. The digested DNA was purified by the PCR purification kit, as the template for IVR. Nanodrop was used to measure the concentration of extracted DNA prior to IVT. Ampliscribe™ T7-Flash™ Transcription Kit was used to generate IVT RNAs as per manufacturer’s instructions. The four canonical (ATP, CTP, GTP and CTP) ribonucleoside triphosphates were supplemented during IVT for producing unmodified RNA oligos. Meanwhile, modified ribonucleoside triphosphates including N1-Methyl-ATP (m1A) and N6-Methyl-ATP (m6A) were supplemented in place of their unmodified counterparts for producing modified RNA oligos. DNAse I was added to the IVT reaction system after incubation for 4 hours at 42 °C to eliminate the residual template DNA. Yielded IVT RNAs were purified using the RNeasy Mini Kit following manufacturer’s instructions. NEB vaccinia capping enzyme was used for the 5’ capping of purified IVT RNAs, with an incubation for 30 min at 37 °C. Following purification with RNAClean XP Beads, the capped IVT RNAs were subjected to polyadenylation tailing. Concentration of capped and polyA-tailed IVT RNAs was determined by Qubit Fluorometer.

### Nanopore Sequencing

RNA nanopore sequencing libraries were built using the ONT Direct RNA Sequencing Kit (SQK-RNA002) following protocol version DRS_9080_v2_revQ_14Aug2019 as per manufacturer’s instructions. Briefly, for each RNA curlcake category, 2 μg of capped and polyA-tailed IVT RNA was subjected to adapter ligation using the NEB T4 DNA Ligase, followed by reverse transcription using the SuperScript III Reverse Transcriptase. After purification using RNAClean XP Beads, yielded RNA:DNA hybrids were ligated to RNA adapters using the NEB T4 DNA Ligase. The concentration of the yielded RNA library was determined using the Qubit fluorometer. The RNA library was mixed with the RNA Running Buffer prior to sequencing on a primed Flongle flowcell. The flowcell version is R9.4.1, and the sequencer is MinION with a Flongle adapter.

### Creating Ground-Truth Sequence Labels for RNA Oligos

We used iterative basecalling to generate ground-truth sequence labels for RNA oligos. During iterative basecalling, we randomly sampled ∼20,000 reads for each modification type (unmodified, m1A, m6A, ac4C, m5C, hm5C, m5U, Psi, m1Psi) and combined them together as a single training dataset. We used Guppy (version 6.0.6+8a98bbc), Taiyaki (version 5.3.0) and Samtools (version 1.16) to perform iterative basecalling. Specifically, we used the Guppy basecalling configuration “rna_r9.4.1_70bps_hac.cfg” for the initial iteration, and the model trained from the previous iteration for the subsequent iteration. The “--disable_qscore_filtering” flag was set to keep “low-quality reads” that are usually artifacts caused by modifications. Samtools functions merge, sort and index with default flags were used to process alignment results generated by Guppy. Taiyaki was used to train basecalling models that are compatible with Guppy. For training data preparation, get_refs_from_sam.py with flag “--reverse”, generate_per_read_params.py with default flags, prepare_mapped_reads.py with the Taiyaki “mLstm_flipflop_model_r941_DNA” checkpoint file, and merge_mappedsignalfiles.py with default flags were used. For training Guppy models, train_flipflop.py with flags “--size 256 --stride 10 --winlen 31” and the model template “mLstm_cat_mod_flipflop.py” were used. For preparing Guppy models, dump_json.py with default flags on the final model checkpoint were used. We performed a total of 4 iterations to guarantee the labeling accuracy, and the comparison between original Guppy and iteratively-optimized basecallers was shown in Figure S6.

### Guppy Basecaller Training and Basecalling Analysis

We trained a total of 18 Guppy basecalling models, including 9 with single modification categories (unmodified, m1A, m6A, ac4C, m5C, hm5C, m5U, Psi, m1Psi), 3 “leave-out” models (trained with all modifications except for ac4C, Psi or m1Psi), as well as 3 “Max” and 3 “Min” models (described in Figure 3B). Training data for individual modification types were prepared from the last iteration of the above-described iterative basecalling process. For “leave-out” models, individual training datasets were combined using the Taiyaki merge_mappedsignalfiles.py function, as above-described. Basecaller models were next trained using the Taiyaki train_flipflop.py and dump_json.py functions, as described in the above section.

Guppy basecalling was performed on independent test datasets: we randomly sampled ∼20,000 reads for each modification type, same as the construction of training datasets. We used flags specified in the configuration file “rna_r9.4.1_70bps_hac.cfg”, except for providing our own models for the Guppy basecalling analysis. Basecalling accuracy was “functionally” evaluated using per-read alignment results, including mappability (the ratio between aligned reads regardless of their SAM flags, as opposed to all the reads), and fractions of CIGAR M (match), X (mismatch), D (deletion) and I (insertion). Specifically, for “leave-out” analyses, we also examined alignment consistency (alignment start and end positions) as opposed to “self” models (model trained with the left-out modification). Alignment analyses were performed with the built-in minimap2 aligner of Guppy, with all the alignment setups set as default.

### Bonito and Dorado Basecaller Training and Basecalling Analysis

We trained a total of 9 Bonito basecalling models, including 9 with single modification categories (unmodified, m1A, m6A, ac4C, m5C, hm5C, m5U, Psi, m1Psi), 3 “leave-out” models (trained with all modifications except for ac4C, Psi or m1Psi). Training data for individual modification types was prepared from the last iteration of the above-described iterative basecalling process. Basecaller models were next trained with the “bonito train” command using the same model architecture as the “rna002_70bps_hac@v3” model. We used the “bonito basecaller” command to basecall test datasets. We used the “bonito export” command to transform the Bonito model to Dorado model and used the “dorado basecaller” command to basecall the same test datasets as in the Guppy analyses were used.

### Representation Space Analysis

Representation space analyses were performed on all possible pairs of basecallers, e.g. above-mentioned single and “leave-out” models and test oligo types, e.g. ac4C, Psi and m1Psi. For a specific pair, e.g. the ac4C “leave-out” model and ac4C test data as shown in Figure 2, we first extracted test sequencing signal chunks that could be approximately corresponded to region curlcake_1_5mers_1:800-820 by the basecaller. Specifically, we scanned consecutive signal chunks of 2,000 data points, and collected ones whose first basecalled nucleotides aligned inside curlcake_1_5mers_1:795-800. We subsequently flowed such chunks through the basecaller encoder neural network, in order to produce a latent feature-by-single chunk matrix in the representation space. With such a matrix, we then performed Principal Component Analysis (PCA), and took the first 50 PCs for the final Uniform Manifold Approximation and Projection (UMAP) visualization^21^.

### Nanopore Sequencing Signal Analysis

We first basecalled nanopore sequencing reads using Bonito (version 0.8.1) to produce move-tables, which track signal chunks corresponding to individual nucleotides (stored as mv tags in the generated bam files). We next executed the Remora (version 3.2.0) pipeline “Reference Region Metric Extraction” (https://github.com/nanoporetech/remora/blob/master/notebooks/metrics_api.ipynb) to extract the per-nucleotide signal features (mean, standard deviation and dwell time). Throughout the analysis, all the Bonito and Remora parameters were set as default.

## Data Availability

The ac4C, m5U and m1Psi RNA oligo nanopore sequencing datasets were downloaded from ENA under the accession number PRJEB67632. The m5C RNA oligo nanopore sequencing dataset was downloaded from NCBI under the BioProject PRJNA563591.

The hm5C RNA oligo nanopore sequencing dataset was downloaded from NCBI under the BioProject PRJNA548268. The Psi RNA oligo nanopore sequencing dataset was downloaded from NCBI under the BioProject PRJNA549001. Resequenced unmodified, and m6A, as well as newly generated m1A nanopore sequencing datasets are available at NCBI under the BioProject PRJNA1050579. Sequence backbones for RNA oligos were provided in the Supplementary Note 1 of ^3^.

## Code Availability

The workflow is available at: https://github.com/wangziyuan66/NanoRL.

## ACKNOWLEDGEMENTS

We thank the University of Arizona High Performance Computing team and the College of Pharmacy Information Technology Group for their support. H.D. is supported by HL166330, the University of Arizona Health Sciences Career Development Award and the University of Arizona Accelerate For Success Award. J.Q. is supported by HL159675, HL152293, AI163753 and DK132251.

## AUTHOR CONTRIBUTIONS

H.D. conceived the idea. Z.W., Z.L. and H.D. performed the analysis. Y.F. performed the experiment. H.H.Z., X.S., N.H., J.Q. and H.D. supervised the project. Z.W., Y.F. and H.D. wrote the manuscript.

