## Supplementary for "Training Data Diversity Enhances the Basecalling of Novel RNA Modification-Induced Nanopore Sequencing Readouts"

### Supplementary Information

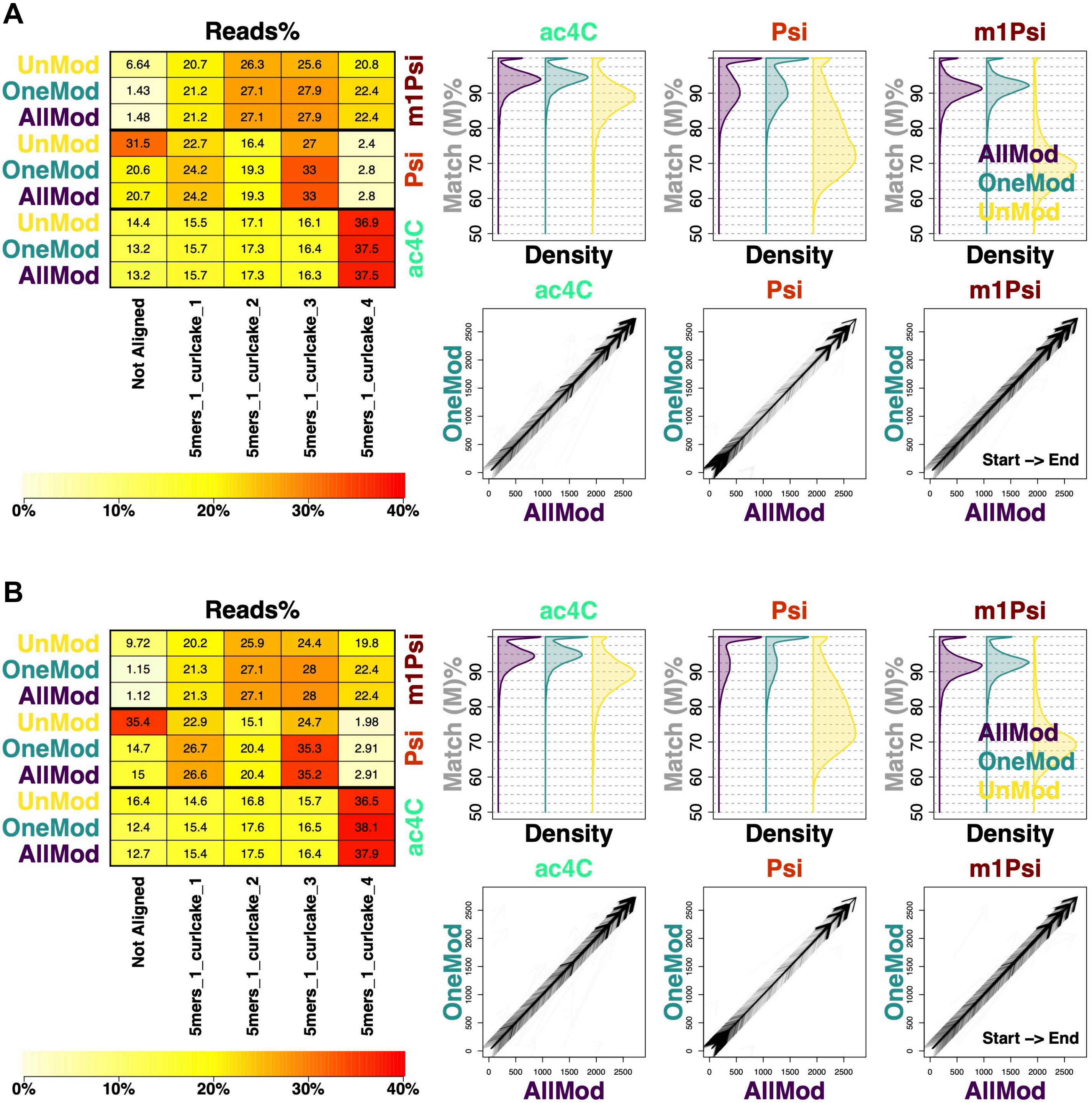
A
Reads%
UnMod
OneMod
AllMod
m1Psi
6.64 20.7 26.3 25.6 20.8
1.43 21.2 27.1 27.9 22.4
1.48 21.2 27.1 27.9 22.4
Psi
31.5 22.7 16.4 27 2.4
20.6 24.2 19.3 33 2.8
20.7 24.2 19.3 33 2.8
ac4C
14.4 15.5 17.1 16.1 36.9
13.2 15.7 17.3 16.4 37.5
13.2 15.7 17.3 16.3 37.5
Not Aligned
5mers_1_curlcake_1
5mers_1_curlcake_2
5mers_1_curlcake_3
5mers_1_curlcake_4
0%
10%
20%
30%
40%
ac4C
Psi
m1Psi
Match (M)%
Density
AllMod
OneMod
UnMod
Start -> End
B
Reads%
m1Psi
9.72 20.2 25.9 24.4 19.8
1.15 21.3 27.1 28 22.4
1.12 21.3 27.1 28 22.4
Psi
35.4 22.9 15.1 24.7 1.98
14.7 26.7 20.4 35.3 2.91
15 26.6 20.4 35.2 2.91
ac4C
16.4 14.6 16.8 15.7 36.5
12.4 15.4 17.6 16.5 38.1
12.7 15.4 17.5 16.4 37.9

**Figure S1. Bonito and Dorado trained with diverse known modifications gain the capability to basecall novel modifications.** The basecalling performance of Bonito and Dorado were presented in (A) and (B), respectively. Same as in Figure 2B-D, the mappability, per-read CIGAR match fraction and alignment consistency were used as quantification metrics. AllMod, the basecaller trained by all the modifications except for the one to be basecalled; OneMod, the basecaller trained with only the modification to be basecalled; UnMod, the basecaller trained by only unmodified reads.

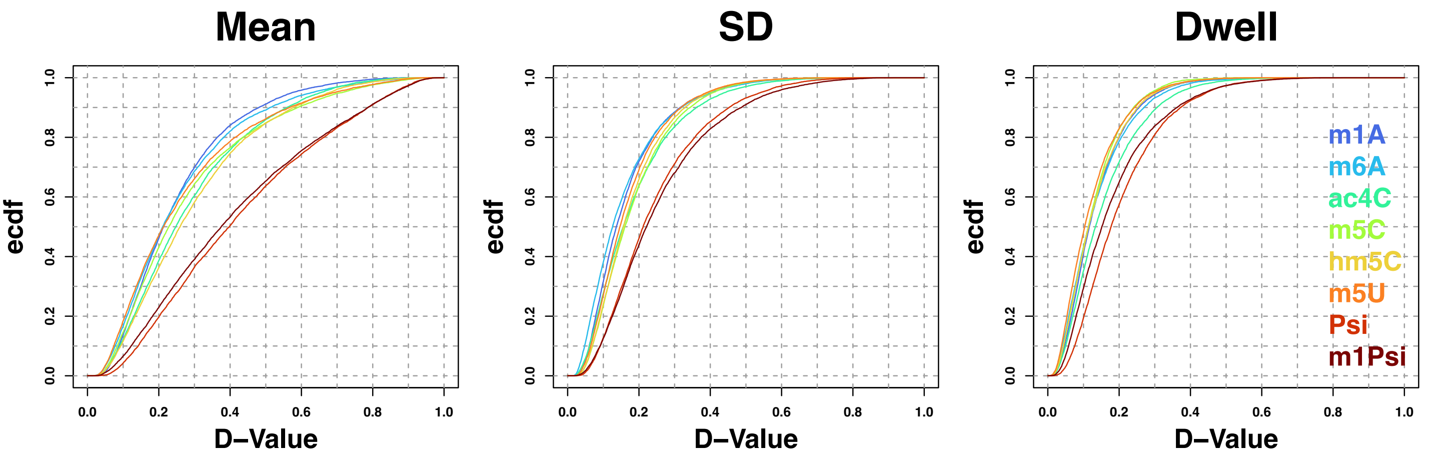
Mean
SD
Dwell
ecdf
D−Value
m1A
m6A
ac4C
m5C
hm5C
m5U
Psi
m1Psi

**Figure S2. Modifications deviate nanopore sequencing signal features.** Mean, SD and Dwell denote the mean, standard deviation and dwell time of Remora signal events. Acronyms denote different modification oligo categories. The KS-test D-values measure Mean, SD and Dwell distribution differences between modified and unmodified oligos. Ecdf denotes the empirical cumulative distribution function.

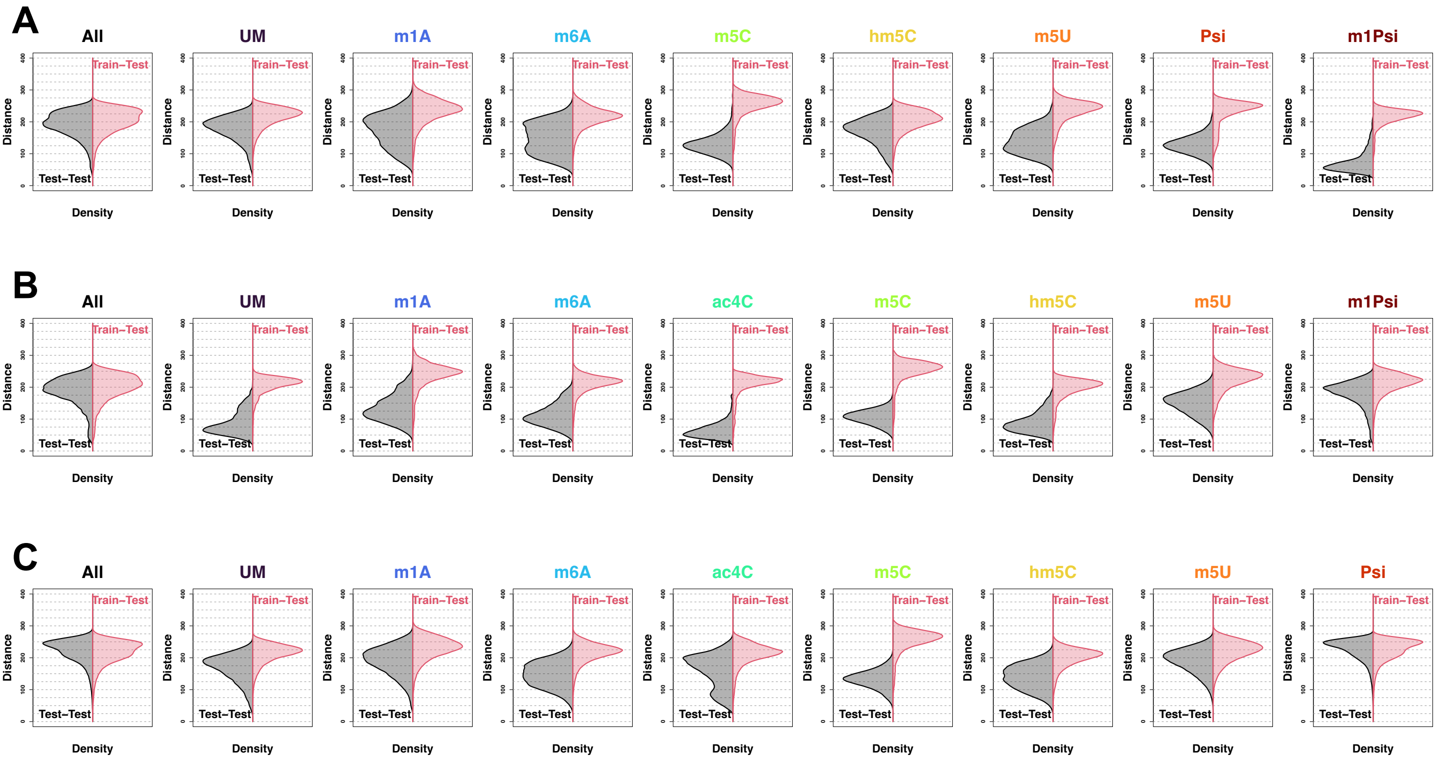

A
All
UM
m1A
m6A
m5C
hm5C
m5U
Psi
m1Psi
B
All
UM
m1A
m6A
ac4C
m5C
hm5C
m5U
m1Psi
C
All
UM
m1A
m6A
ac4C
m5C
hm5C
m5U
Psi
Train−Test
Test−Test
Distance
Density

**Figure S3. Quantifying the read-level similarity in the representation space.** Reads were projected into the representation space and then subjected to PCA. Within the PC space, Euclidean distance was calculated to quantify the read-level similarity. Analyses for ac4C, Psi and m1Psi test reads were presented in (A), (B) and (C), respectively. All, the jointly-trained basecaller by all the oligo groups except for the test; other acronyms denote individually-trained basecallers. Train-Test and Test-Test represent the distance between training and test reads and among test reads, respectively. We took Test-Test groups as baselines to quantify the Train-Test similarity.

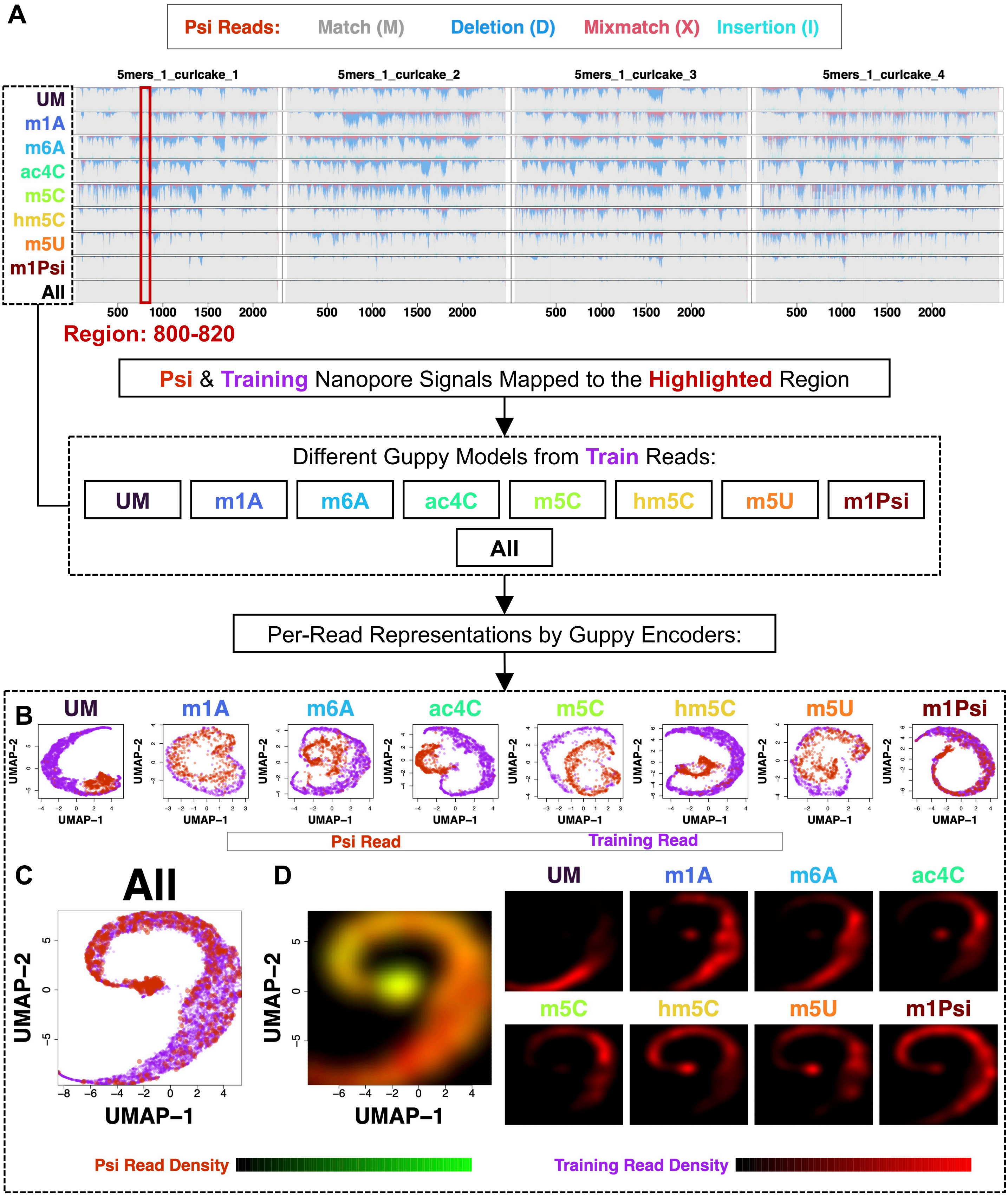
A
Psi Reads: Match (M) Deletion (D) Mixmatch (X) Insertion (I)
5mers_1_curlcake_1
5mers_1_curlcake_2
5mers_1_curlcake_3
5mers_1_curlcake_4
UM
m1A
m6A
ac4C
m5C
hm5C
m5U
m1Psi
All
Region: 800-820
Psi & Training Nanopore Signals Mapped to the Highlighted Region
Different Guppy Models from Train Reads:
UM m1A m6A ac4C m5C hm5C m5U m1Psi
All
Per-Read Representations by Guppy Encoders:
B
UMAP-1
UMAP-2
Psi Read
Training Read
C
All
D
Psi Read Density
Training Read Density

**Figure S4. A generalizable representation space produced from diverse training modifications facilitates the basecalling of the out-of-sample Psi modification.** (A) Performance of individually and jointly-trained basecallers on Psi reads was visualized with the genome viewer graph, which shows per-nucleotide CIGAR fractions. All, the jointly-trained basecaller by all the oligo types except for Psi; other acronyms denote individually-trained basecallers. For individually (B) and jointly-trained (C) basecallers, read fragments mapped to the boxed region were first converted as representation vectors with different basecaller encoders, then visualized by a UMAP plot. Train denotes reads used for training the corresponding basecaller. (D) Spatial distributions of different oligo types in the UMAP space as shown in (C). Black-to-green and red palette denotes Psi and training reads, respectively.

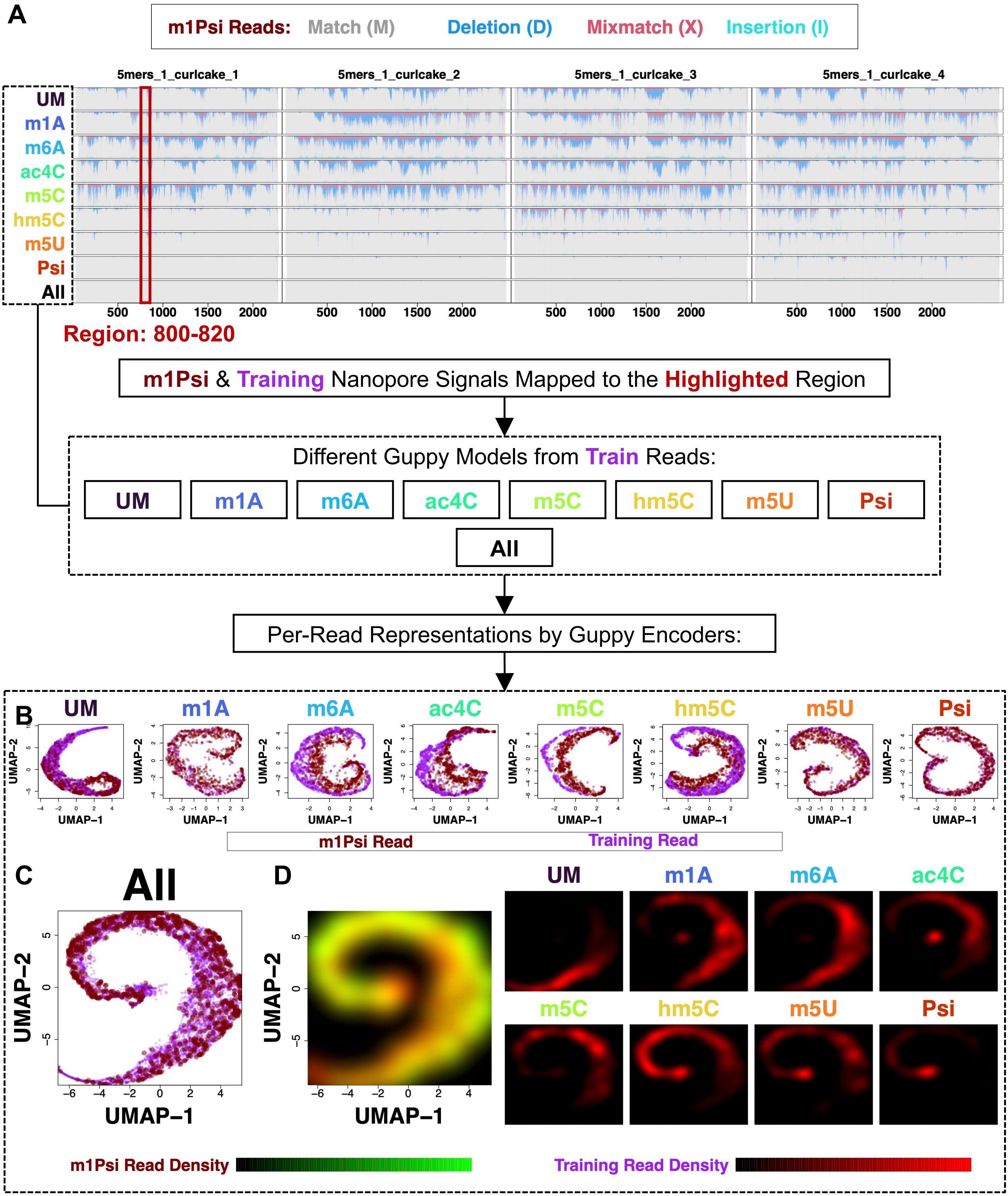
A
m1Psi Reads: Match (M) Deletion (D) Mixmatch (X) Insertion (I)
5mers_1_curlcake_1
5mers_1_curlcake_2
5mers_1_curlcake_3
5mers_1_curlcake_4
UM
m1A
m6A
ac4C
m5C
hm5C
m5U
Psi
All
500 1000 1500 2000
Region: 800-820
m1Psi & Training Nanopore Signals Mapped to the Highlighted Region
Different Guppy Models from Train Reads:
UM m1A m6A ac4C m5C hm5C m5U Psi
All
Per-Read Representations by Guppy Encoders:
B
UMAP–1
UMAP–2
m1Psi Read
Training Read
C
All
D
m1Psi Read Density
Training Read Density

**Figure S5. A generalizable representation space produced from diverse training modifications facilitates the basecalling of the out-of-sample m1Psi modification.** (A) Performance of individually and jointly-trained basecallers on m1Psi reads was visualized with the genome viewer graph, which shows per-nucleotide CIGAR fractions. All, the jointly-trained basecaller by all the oligo types except for m1Psi; other acronyms denote individually-trained basecallers. For individually (B) and jointly-trained (C) basecallers, read fragments mapped to the boxed region were first converted as representation vectors with different basecaller encoders, then visualized by a UMAP plot. Train denotes reads used for training the corresponding basecaller. (D) Spatial distributions of different oligo types in the UMAP space as shown in (C). Black-to-green and red palette denote m1Psi and training reads, respectively.

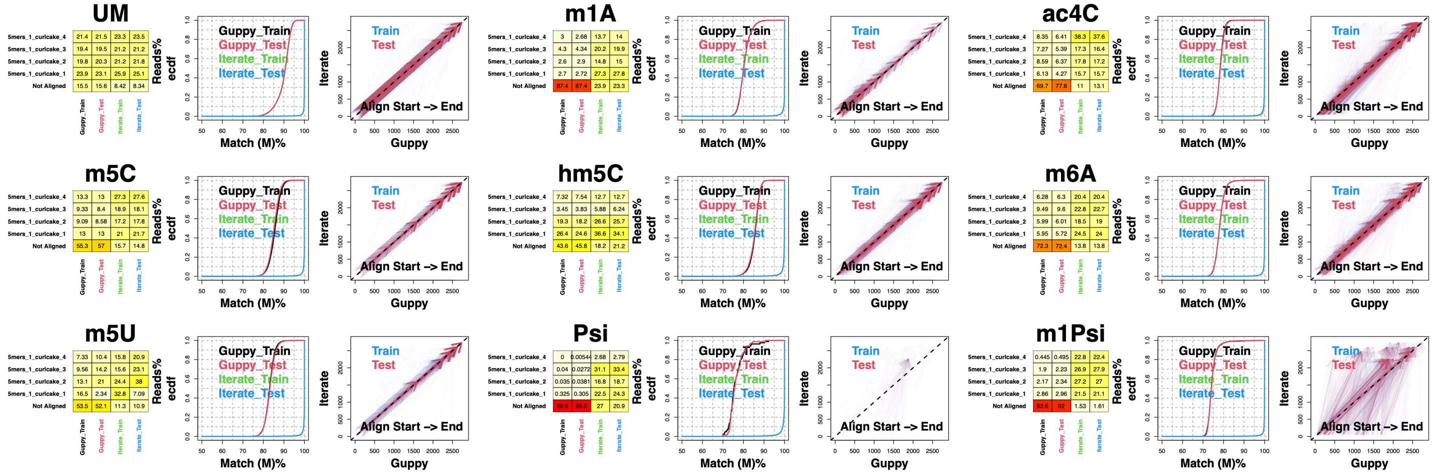
UM
5mers_1_curlcake_4 21.4 21.5 23.3 23.5
5mers_1_curlcake_3 19.4 19.5 21.2 21.2
5mers_1_curlcake_2 19.8 20.3 21.2 21.8
5mers_1_curlcake_1 23.9 23.1 25.9 25.1
Not Aligned 15.5 15.6 8.42 8.34
m1A
5mers_1_curlcake_4 3 2.68 13.7 14
5mers_1_curlcake_3 4.3 4.34 20.2 19.9
5mers_1_curlcake_2 2.6 2.9 14.8 15
5mers_1_curlcake_1 2.7 2.72 27.3 27.8
Not Aligned 87.4 87.4 23.9 23.3
ac4C
5mers_1_curlcake_4 8.35 6.41 38.3 37.6
5mers_1_curlcake_3 7.27 5.39 17.3 16.4
5mers_1_curlcake_2 8.59 6.37 17.8 17.2
5mers_1_curlcake_1 6.13 4.27 15.7 15.7
Not Aligned 69.7 77.6 11 13.1
m5C
5mers_1_curlcake_4 13.3 13 27.3 27.6
5mers_1_curlcake_3 9.33 8.4 18.9 18.1
5mers_1_curlcake_2 9.09 8.58 17.2 17.8
5mers_1_curlcake_1 13 13 21 21.7
Not Aligned 55.3 57 15.7 14.8
hm5C
5mers_1_curlcake_4 7.32 7.54 12.7 12.7
5mers_1_curlcake_3 3.45 3.83 5.88 6.24
5mers_1_curlcake_2 19.3 18.2 26.6 25.7
5mers_1_curlcake_1 26.4 24.6 36.6 34.1
Not Aligned 43.6 45.8 18.2 21.2
m6A
5mers_1_curlcake_4 6.28 6.3 20.4 20.4
5mers_1_curlcake_3 9.49 9.6 22.8 22.7
5mers_1_curlcake_2 5.99 6.01 18.5 19
5mers_1_curlcake_1 5.95 5.72 24.5 24
Not Aligned 72.3 72.4 13.8 13.8
m5U
5mers_1_curlcake_4 7.33 10.4 15.8 20.9
5mers_1_curlcake_3 9.56 14.2 15.6 23.1
5mers_1_curlcake_2 13.1 21 24.4 38
5mers_1_curlcake_1 16.5 2.34 32.8 7.09
Not Aligned 53.5 52.1 11.3 10.9
Psi
5mers_1_curlcake_4 0 0.00544 2.68 2.79
5mers_1_curlcake_3 0.04 0.0272 31.1 33.4
5mers_1_curlcake_2 0.035 0.0381 16.8 18.7
5mers_1_curlcake_1 0.325 0.305 22.5 24.3
Not Aligned 99.6 99.6 27 20.9
m1Psi
5mers_1_curlcake_4 0.445 0.495 22.8 22.4
5mers_1_curlcake_3 1.9 2.23 26.9 27.9
5mers_1_curlcake_2 2.17 2.34 27.2 27
5mers_1_curlcake_1 2.86 2.96 21.5 21.1
Not Aligned 92.6 92 1.53 1.61
Guppy_Train
Guppy_Test
Iterate_Train
Iterate_Test
Reads%
ecdf
Match (M)%
Train
Test
Iterate
Guppy
Align Start –> End

**Figure S6. The iterative basecalling of RNA oligos.** Guppy and iterate denote Guppy and 4th iteration (final iteration) basecallers, respectively. Train and test denote training and the independent test datasets, respectively. UM denotes unmodified reads. Match denotes the CIGAR category M. Ecdf, empirical cumulative distribution function. Start and End denote the alignment direction.
